# Identification and verification of 3 key genes associated with survival and prognosis of patients with colon adenocarcinoma via integrated bioinformatics analysis

**DOI:** 10.1101/2020.06.09.142042

**Authors:** Yong Liu, Chao Li, Lijin Dong, Ping Li, Xuewei Chen, Rong Fan

**Author notes:** Correspondence: Xuewei Chen, Department of Operational Medicine, Tianjin Institute of Environmental and Operational Medicine, Tianjin, 300050, PR China, E-mail address, Rong Fan, Central Lab, Tianjin XiQing hospital, Tianjin 300380, PR China. These authors contributed equally to this work (Yong Liu, Chao Li).

## Abstract

**Background:** Colorectal cancer (CRC) is the third most lethal malignancy in the world, wherein colon adenocarcinoma (COAD) is the most prevalent type of CRC. Exploring biomarkers is important for the diagnosis, treatment, and prevention of COAD.

**Methods:** We used GEO2R and Venn online software for differential gene screening analysis. Hub genes were screened via STRING and Cytoscape, following Gene Ontology and KEGG enrichment analysis. Finally, survival analysis and expression validation were performed via UALCAN online software, real-time PCR and immunohistochemistry.

**Results:** In this study, we screened 323 common differentially expressed genes from four GSE datasets. Furthermore, four hub genes were selected for survival correlation analysis and expression level verification, three of which were shown to be statistically significant.

**Conclusion:** Our study suggests that SERPINE1, SPP1 and TIMP1 may be biomarkers closely related to the prognosis of CRC patients.

## 1. Introduction

Colorectal cancer (CRC) originates in the colon or rectum [1], and typical symptoms include alternating diarrhea, constipation, mucus, bloody stools, as well as weight loss [2]. According to annual reports on global cancer statistics, nearly 1.4 million newly diagnosed cases and 700,000 deaths occur worldwide due to CRC [3]. The five year survival rate of patients with CRC at its early stage is 90 percent, while those at later stages show a rate of no more than 12 percent [4, 5]. Colon adenocarcinoma (COAD) is one of the most prevalent types of CRC, the incidence and mortality of which was 10.2% and 9.2%, respectively, in 2018 [6, 7]. Therefore, it is very important to select and identify the specific biomarkers of COAD for its early diagnosis, development of an efficient treatment strategy, and assessment of patient prognosis.

Gene microarray technology has been extensively applied in several fields of medicine and biology. This high-throughput technology can effortlessly acquire gene expression profile information, which is very useful for exploring various molecular mechanisms [8]. Recently, integrated bioinformatics analysis has also been widely applied; it includes mathematics and biology, which makes it possible to analyze large-scale microarray data. In addition, it is advantageous as an efficient method for the systematical screening of tumor-associated genes [9, 10].

In this study, we screened the common differentially expressed genes (DEGs) in 4 GSE datasets from the Gene Expression Omnibus (GEO) repository and constructed a protein-protein interaction (PPI) network. Via integrated bioinformatics analysis, we identified the 4 hub genes associated with survival of COAD patients and verified the mRNA expression levels in CRC cells. Finally, our results showed that SPP1, SERPINE1 and TIMP1 could be potential biomarkers for prognostics of COAD in patients.

## 2. Materials and Methods

### 2.1 Download of microarray data

The high-throughput gene expression profiles of normal colorectal tissues and COAD tissues were extracted from the GEO database. The independent datasets of GSE37364, GSE41328, GSE81558 and GSE110225 were selected which included 14 COAD tissues and 37 normal tissues, 10 COAD tissues and 10 normal tissues, 23 COAD tissues and 9 normal tissues, and 17 COAD tissues and 17 normal tissues, respectively.

### 2.2 Identification of DEGs

The DEGs between normal colorectal tissues and COAD tissues were screened via the GEO2R tool. We limited the cut-off criterion of | logFC | ≥ 1 to be statistically significant. Then, the DEGs in the four datasets were screened out using online Venn software. The DEGs with logFC ≥1 were considered as upregulated genes, while DEGs with logFC ≤-1 were considered as downregulated genes.

### 2.3 Enrichment analysis via GO and KEGG pathway

In order to characterize the functional roles of the DEGs, we used DAVID (version 6.8 [11]) database for gene ontology and signal pathway enrichment analysis, which comprised biological process (BP), molecular function (MF) and cellular component (CC), and KEGG pathways analysis.

### 2.4 Construction of PPI network and analysis of module

The PPI network was built via the STRING [12] database to uncover the relationships of DEGs, based on a minimum required interaction score of 0.4. We used Cytoscape (version 3.7.1) [13] software to complete the analysis and visualization of the PPI network. In addition, the remarkable modules of PPI network were selected via the MCODE app (node score cutoff=0.2, degree cutoff=2, max. Depth=100, and k-score=2).

### 2.5 Survival and RNA expression analysis of hub genes

UALCAN [14] was used for analyzing the relationship between hub genes expression and survival of patients with COAD, as it is a good tool for analyzing transcriptome data of cancers from The Cancer Genome Atlas (TCGA). Simultaneously, UALCAN was used to analyze RNA expression data based on the TCGA. *P*-value <0.05 was considered statistically significant.

### 2.6 Cell lines and culture

The normal colon epithelial cell line, NCM460, and colon adenocarcinoma cell lines, HCT116, HCA-7, HT-29, and SW480 were maintained by our laboratory. And all cells were cultured in a 5% CO_2_ humidified incubator in Dulbecco’s Modified Eagle’s medium (Hyclone) containing 10% fetal bovine serum (FBS, Gibco) at 37°C.

### 2.7 Quantitative real-time Polymerase Chain Reaction (PCR)

Total RNA from cell cultures was collected by using Total RNA Extraction Kit (Solarbio Life Sciences, China). The specific primers were designed by using Primer Express 3.0 software. Each sample was used in triplicate for the qPCR using PowerUp™ SYBR™ Green Master Mix (Thermo Fisher Scientific, USA) on Applied Biosystems™ StepOnePlus instruments, and analyzed with the StepOne software, using GAPDH as a control. The primers used in this study are designed as follows: TIMP1: Forward:5’-GGGACACCAGAAGTCAACCAG-3’; Reverse:5’-CAATGAGAAACTCCTCGCTGC-3’; CXCL2: Forward: 5’-CCAAACCGAAGTCATAGCCAC-3’; Reverse: 5’-TGCCATTTTTCAGCATCTTTTC-3’; SPP1: Forward: 5’-TGAGCATTCCGATGTGATTGAT-3’; Reverse: 5’-TTATCTTCTTCCTTACTTTTGGGGT-3’; SERPINE1: Forward: 5’-CTGGGTGAAGACACACACAAAAG-3’; Reverse: 5’-CACAGAGACAGTGCTGCCGT-3’; GAPDH: Forward: 5’-TCAAGGGCATCCTGGGCTACAC-3’; Reverse: 5’-TCAAAGGTGGAGGAGTGGGTGTC-3’.

### 2.8 Tissue samples and immunohistochemistry (IHC)

Human colon adenocarcinoma tissue microarray sections (HColA160CS01) were obtained from Shanghai Outdo Biotech Co. Ltd. (Shanghai, China). The tissue samples were procured from 80 patients with colorectal cancer. Every specimen was provided with cancer tissue and adjacent-carcinoma tissue. The two-step EnVision method was used to perform immune histochemical experiments, along with different primary antibodies against CXCL2 (1:50), SERPINE1 (1:50), TIMP1 (1:50) and SPP1 (1:50). The study was approved by the Ethics Committee of Shanghai Outdo Biotech Company (Shanghai, China). The slides were analyzed using the Image-Pro PLUS software program (Media Cybernetics, Inc. USA).

### 2.9 Statistical analysis

Statistical analysis was performed using SPSS 22.0 and GraphPad Prism 8.0 software. Results were presented as the mean ± standard deviation. The statistically significant changes were indicated with asterisks and the *P*-values were calculated via a Student’s t-test, wherein *, **, and *** represented *P*< 0.05, *P*< 0.01 and *P*< 0.001, respectively.

## 3. Results

### 3.1 Identification of DEGs in COAD

In this study, we selected and download 4 GEO datasets and extracted DEGs on the basis of the cut-off criteria. This study covered 64 COAD and 73 normal colorectal tissues in total. The results showed that 2729, 957, 1864, and 617 genes were down-regulated while 1917, 770, 1282, and 469 genes were upregulated in GSE37364, 41328, 81558, and 110250, respectively. By using the Venn diagram software, we detected a total of 323 common DEGs in the COAD samples, composed of 111 up-regulated genes and 212 down-regulated genes (Table 1 & Figure 1).

**Figure 1.**
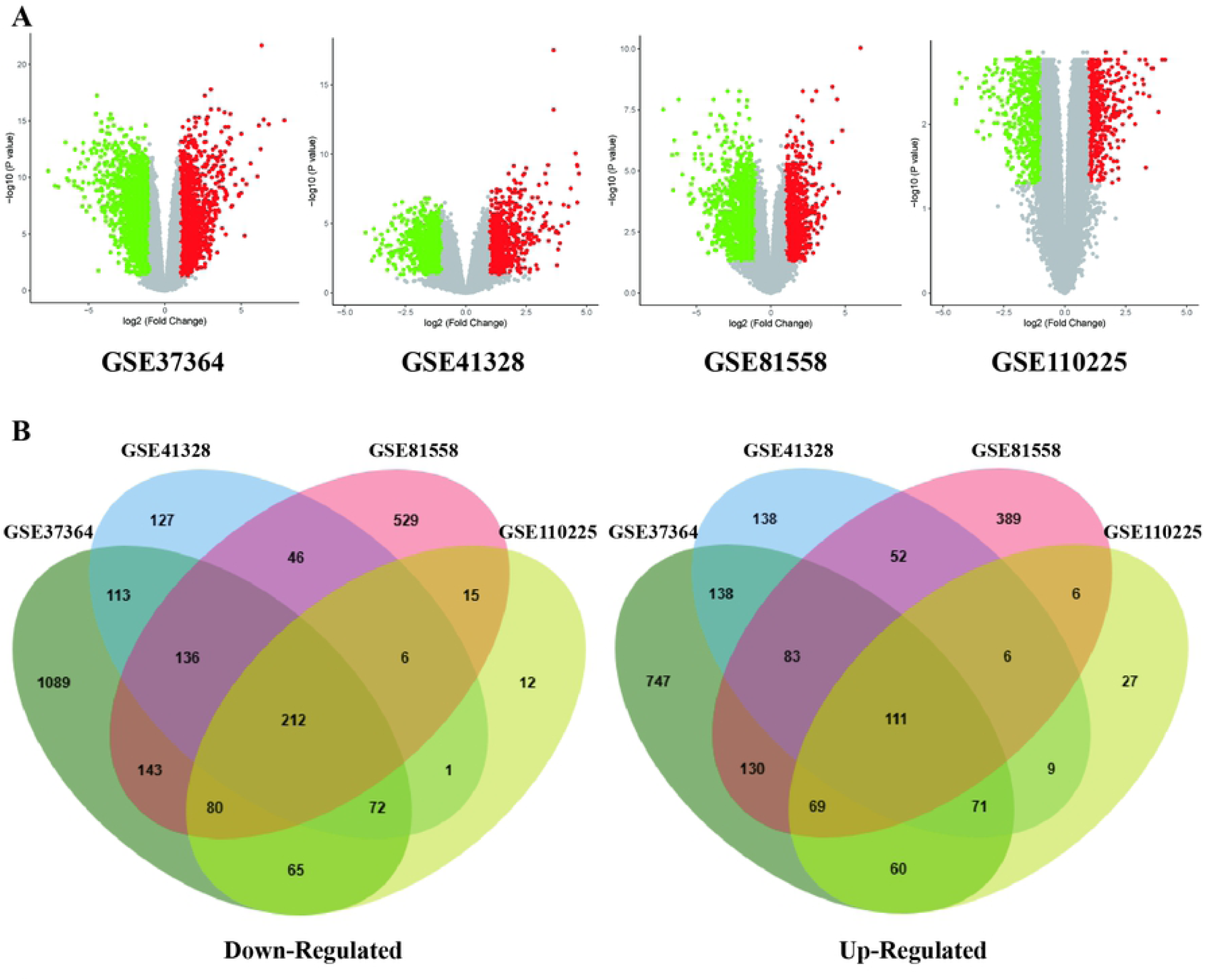
Selection of 323 common DEGs from the four datasets (GSE37364, GSE41328, GSE81558, and GSE110225). A. Volcano plot of DEGs from the four datasets; B. On the left, 136 DEGs are downregulated (logFC≤1); On the right, 198 DEGs are upregulated (logFC≥1).

### 3.2 GO and signaling pathway enrichment analysis

We analyzed the DEGs via the DAVID software and identified 125 significant enrichments terms, including BP (71), MF (24), and CC (26). With regard to BP, DEGs were seen to be significantly enriched during cell adhesion, negative regulation of cell proliferation, angiogenesis, cytoskeleton organization, steroid hormone mediated signaling pathway, inflammatory response, negative regulation of protein kinase activity, and so on. Whereas, for MF, DEGs were significantly enriched during chemokine activity, hormone activity, CXCR chemokine receptor binding, receptor binding, calcium ion binding, structural constituent of cytoskeleton, protein kinase inhibitor activity, oxidoreductase activity, etc. With regard to CC, DEGs were significantly enriched in extracellular space, apical plasma membrane, extracellular exosome, extracellular region, cell surface, plasma membrane, basal plasma membrane, and so forth. The top 20 GO terms are depicted in Figure 2 A-C. Furthermore, KEGG pathway analysis indicated that the DEGs were enriched in 11 pathways including nitrogen metabolism, bile secretion, proximal tubule bicarbonate reclamation, cytokine-cytokine receptor interaction, aldosterone-regulated sodium reabsorption, mineral absorption, PI3K-Akt signaling pathway, PPAR signaling pathway, etc. (Figure 2D).

**Figure 2.**
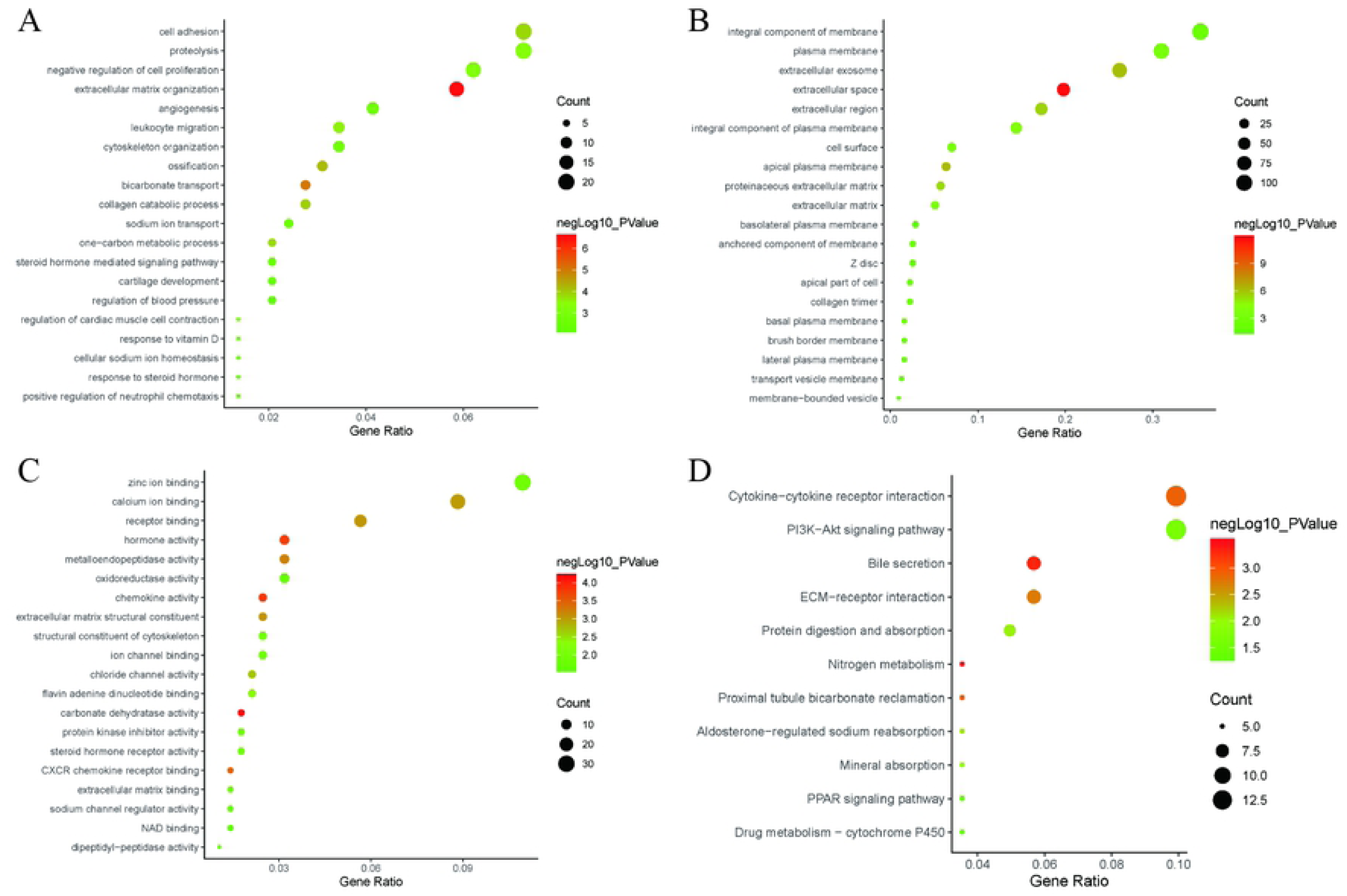
The top 20 gene ontology (GO) and significantly enriched KEGG pathways. A. Biological process (BP); B. Cellular components (CC); C. Molecular functions (MF); D. KEGG pathways.

### 3.3 Construction of PPI network and module analysis

Via the STRING and Cytoscape analysis, we drew the PPI network complex constructed with 268 nodes and 709 edges, consisting of 97 downregulated and 171 upregulated genes. Then we conducted further analyses by applying Cytoscape MCODE plus, and found 32 central nodes that included 20 upregulated genes and 12 downregulated genes (Figure 3, 4 & Table 2).

**Figure 3.**
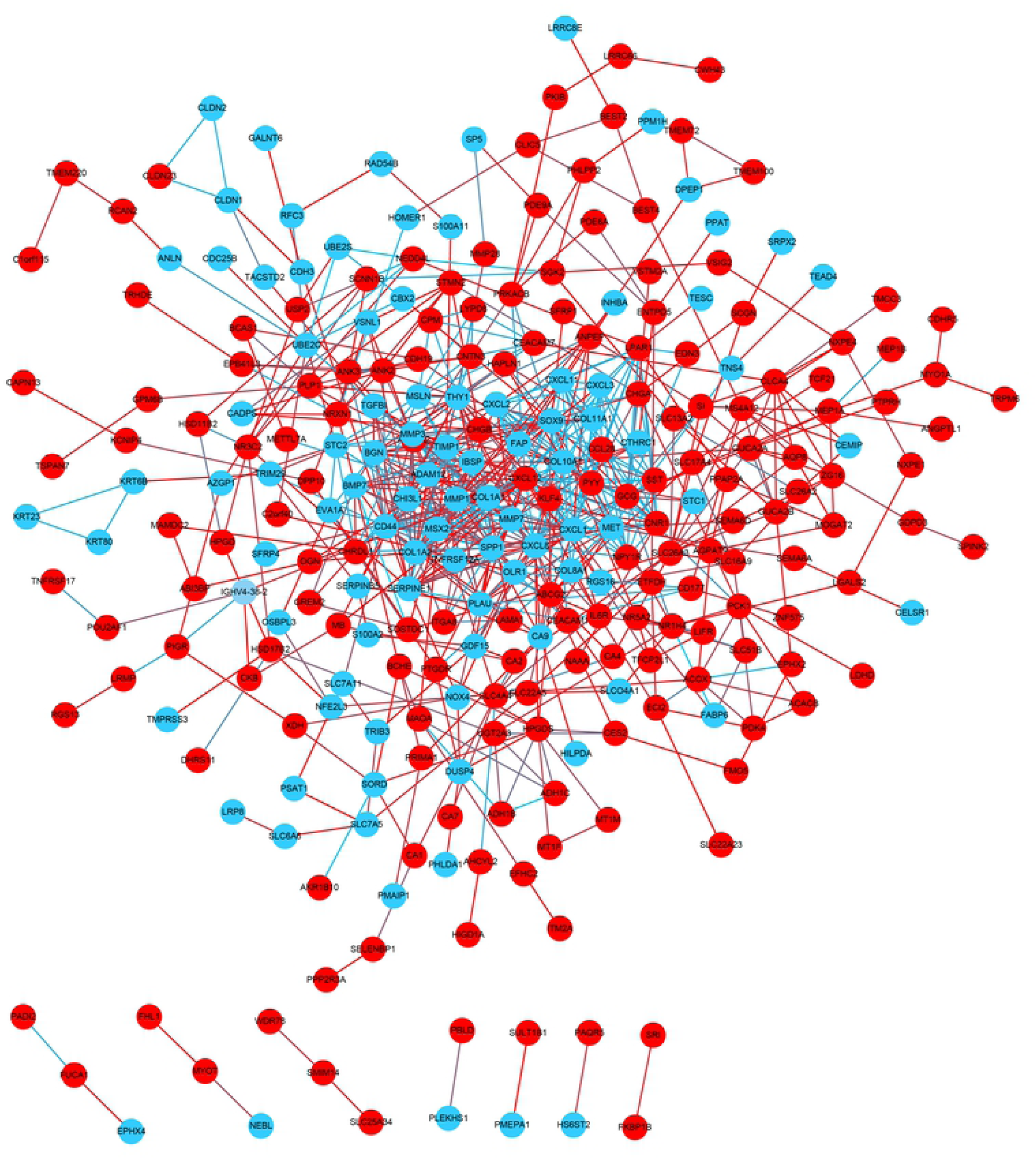
PPI network constructed via STRING and Cytoscape. Each node indicates a protein module; the edges represent proteins interaction; blue circles denote upregulated DEGs while red circles represent downregulated DEGs.

**Figure 4.**
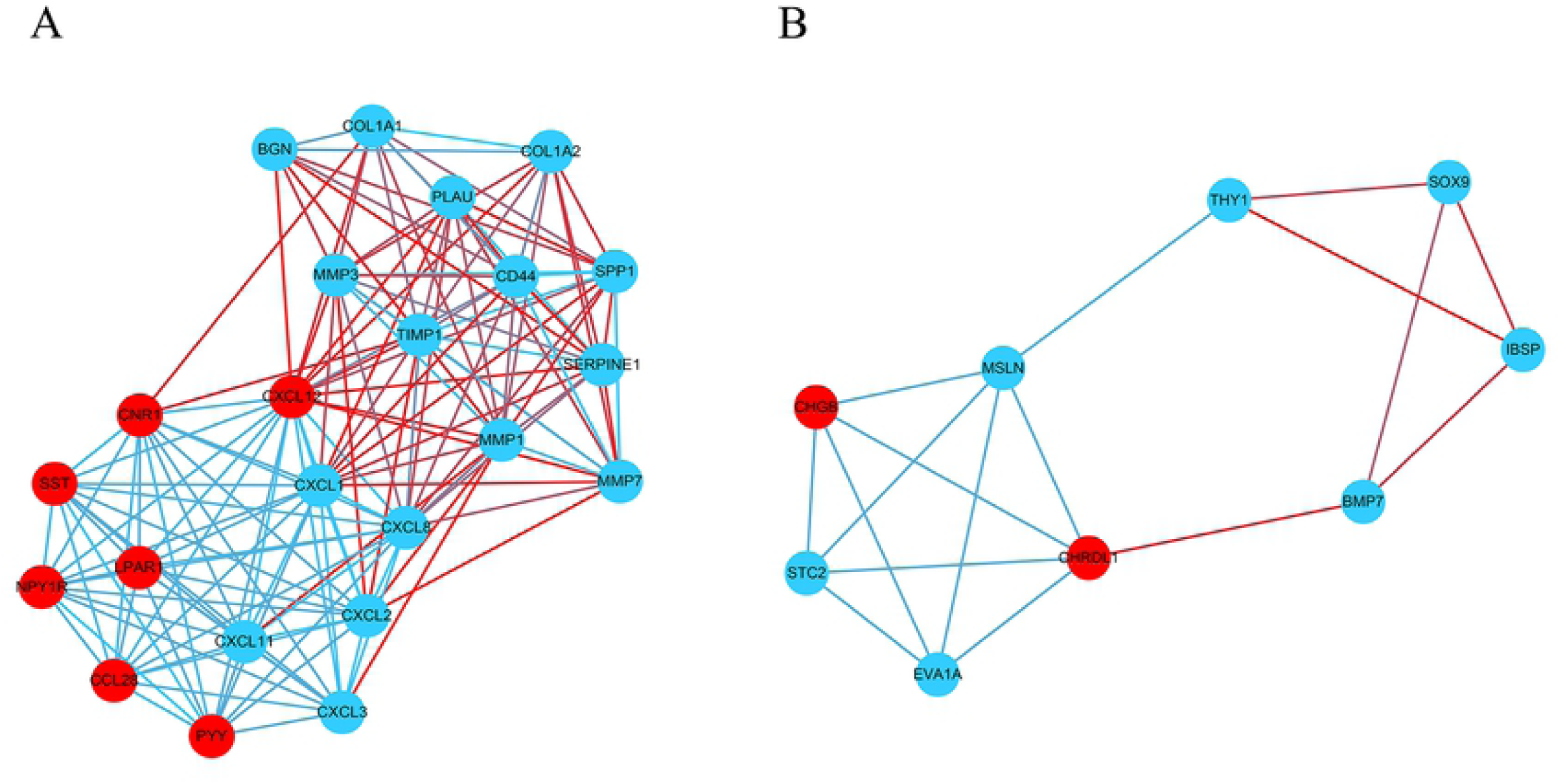
Module analysis by MCODE in the Cytoscape software. A. module1; B. module2. (Blue circles signify upregulated DEGs and red circles denoted downregulated DEGs)

### 3.4 Selection of hub genes and validation of the expression levels

We used UALCAN to identify the survival data of 32 central genes and found that the expression levels of 4 hub genes, namely *CXCL2, SERPINE1, SPP1*, and *TIMP1*, were remarkably related to the survival of COAD patients (Figure 5). Furthermore, we validated the expression level of these 4 hub genes in COAD and normal tissues via UALCAN. Results showed that expression levels of the 4 genes were significantly increased in the COAD samples as compared to the normal samples (Figure 6). Furthermore, we verified the mRNA levels of the four hub genes in NCM460, HCT116, HCA-7, HT-29, and SW480 cells. The real-time PCR results showed that the expression levels of SPP1, TIMP1, and SERPINE1 mRNAs were significantly increased in the HCT116, HCA-7, HT-29, and SW480 cell lines as compared to the NCM460 cell line (Figure 7). Moreover, CXCL2 expression was higher in the HCA-7, HT-29, and SW480 cells, but not in the HCT116 cells when compared to NCM460. In addition, we also analyzed the protein expressions of the four hub genes by using tissue microarray. The results indicated that SPP1, TIMP1, and SERPINE1 expression appeared to be remarkably higher in colorectal cancer tissues than in the adjacent tissues, while the expression of CXCL2 was not statistically different between cancer tissues and adjacent tissues (Figure 8).

**Figure 5.**
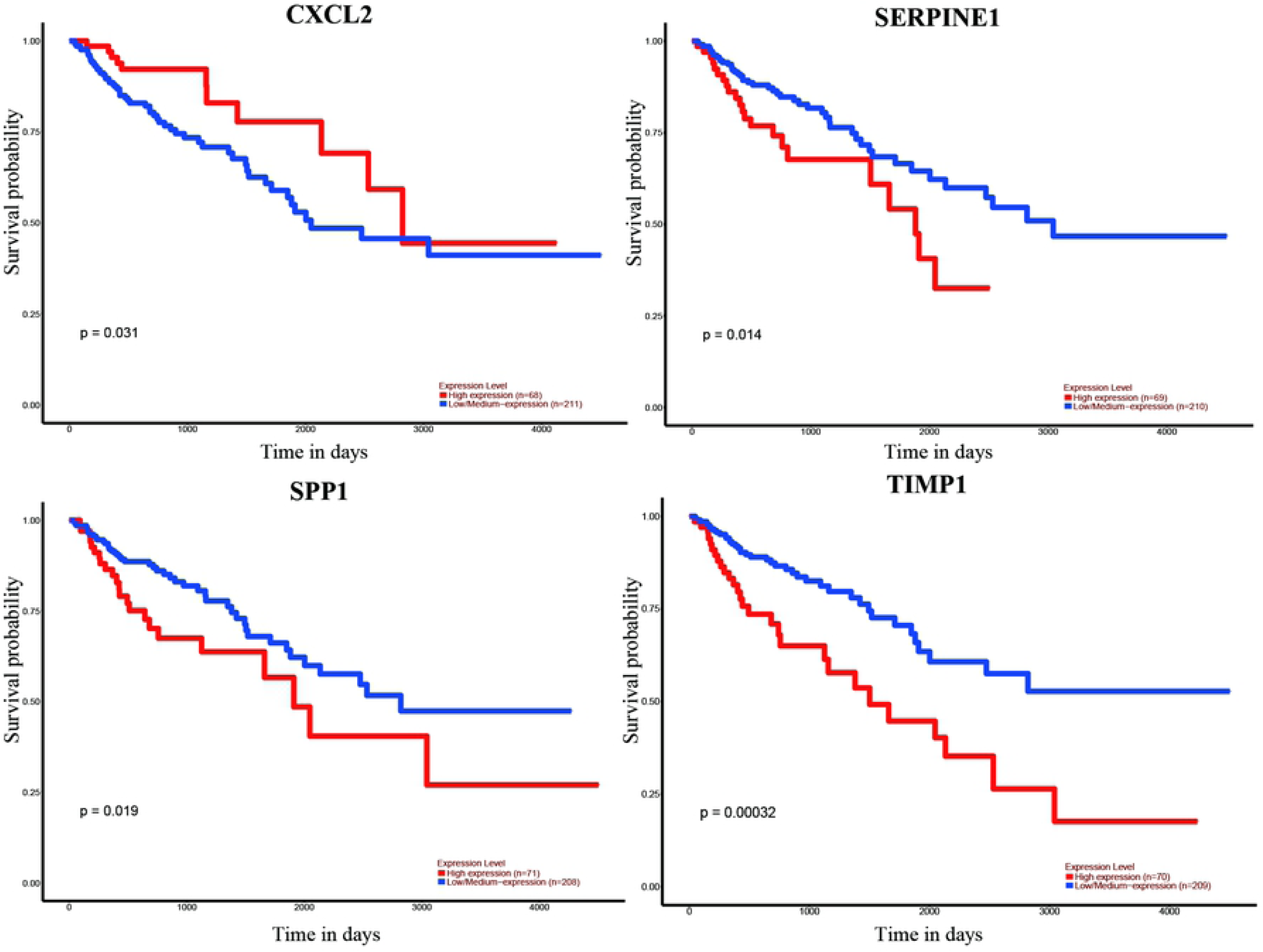
Correlation between the 4 hub genes expression and the overall survival of COAD patients, via UALCAN analysis.

**Figure 6.**
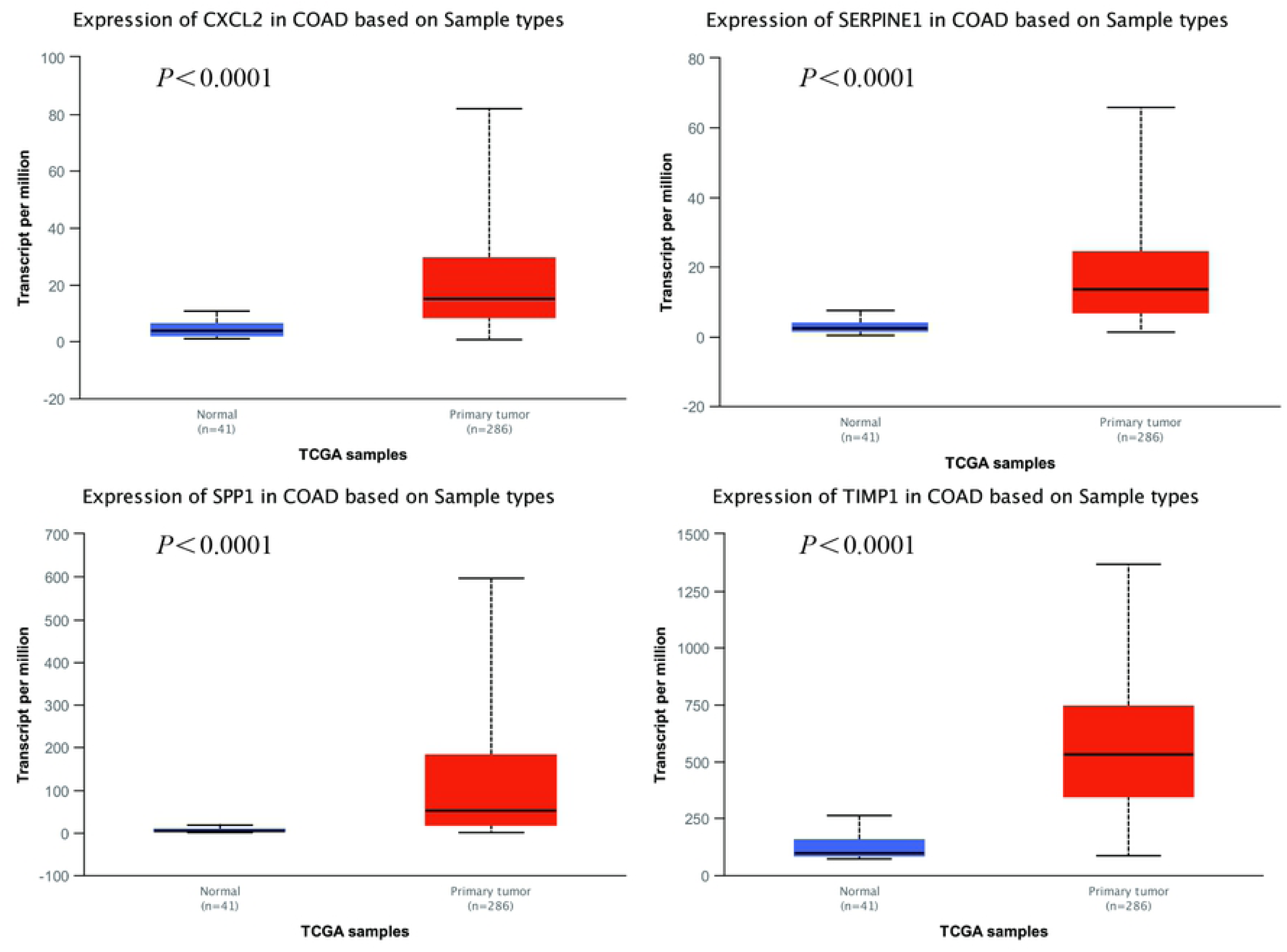
Analysis of the 4 hub genes expression levels via UALCAN analysis.

**Figure 7.**
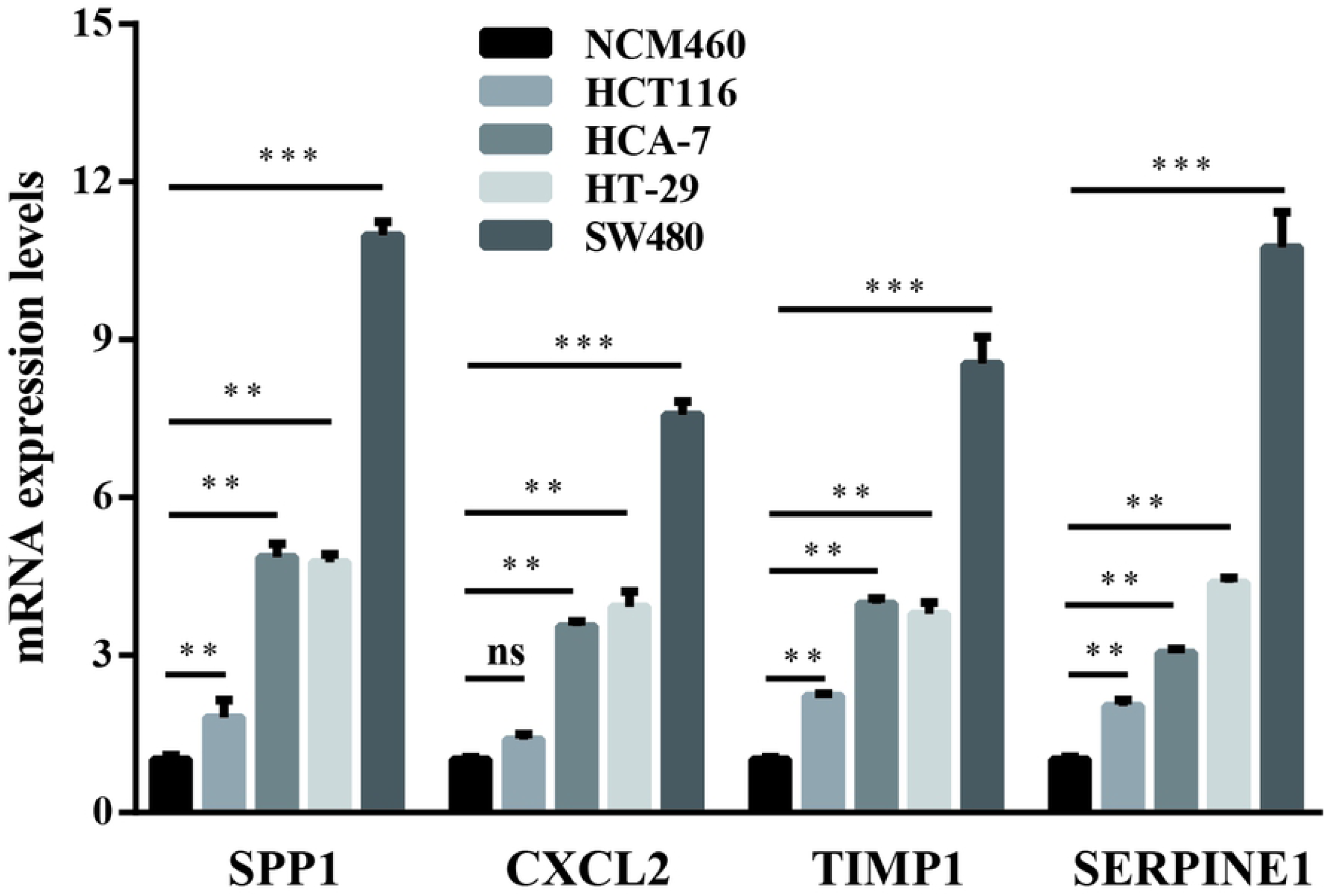
Relative mRNA levels of the 4 hub genes in HCT116, HCA-7, HT-29, and SW480 cells compared to NCM460 cells, via real-time PCR. (** represents *P*<0.01, *** represents *P*<0.001, ns represents no significance)

**Figure 8.**
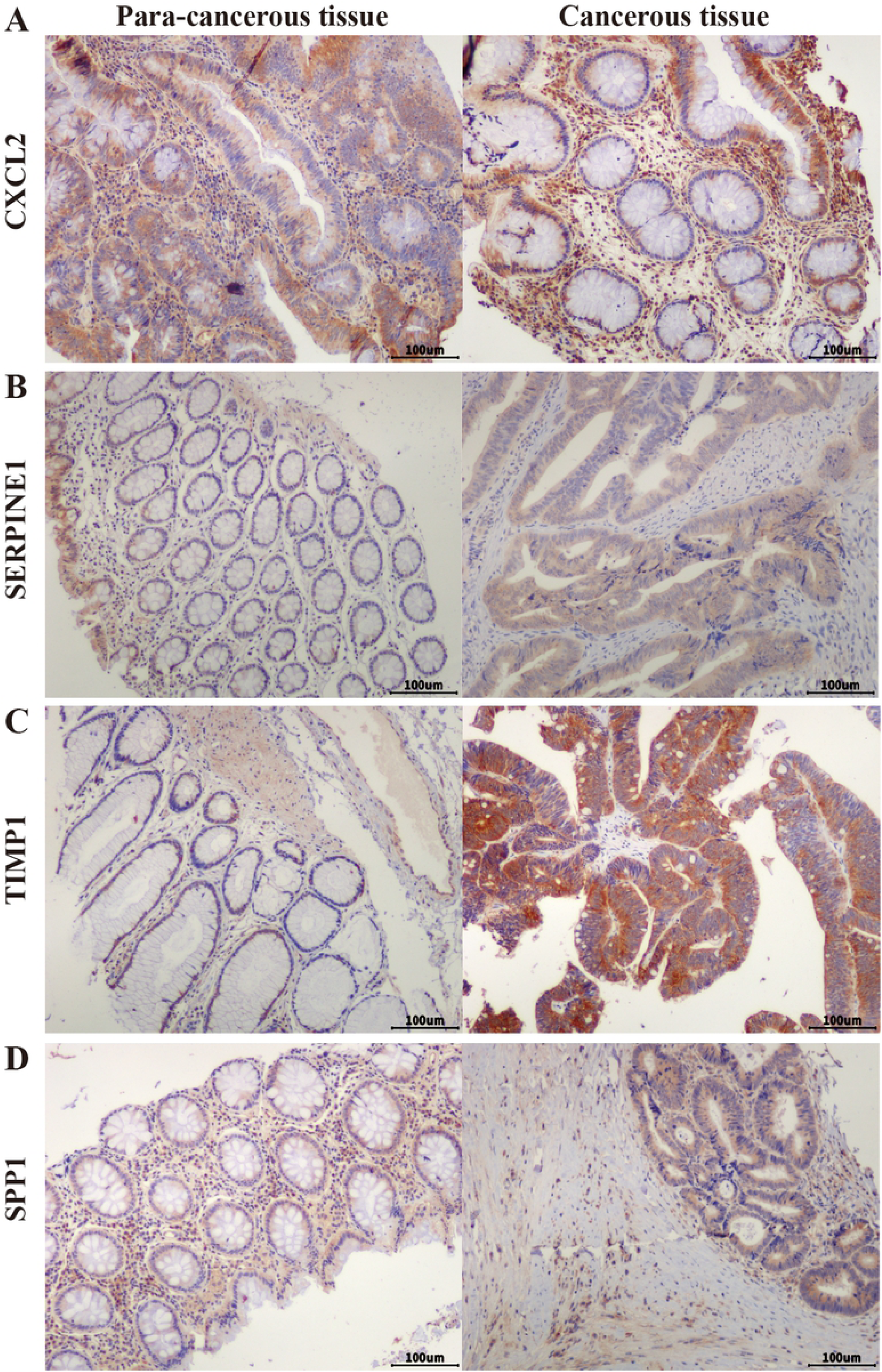
Expression of CXCL2 (A), SERPINE1 (B), TIMP1 (C), and SPP1 (D) in colorectal adenocarcinoma tissue and adjacent tissue in tissue microarrays. (Scale bar: 100μm)

## 4. Discussion

CRC is the third main cause of death from malignant cancers worldwide, and the fourth most common cancer in China [15, 16]. One of the most common types of CRC is COAD, which originates in the superficial glandular epithelial cells [6]. Previous studies have indicated that the mechanism behind the tumorigenesis and development of COAD is a complicated process, including chromosomal instability, dysregulation of oncogenes and tumor suppressor, epigenetic alterations, metabolic alterations, abnormal immune response, etc. [17].

In this study, we screened DEGs via GEO2R, and selected 111 up-regulated genes and 212 down-regulated genes found in COAD tissues. Then we analyzed their enrichment in GO and signaling pathways and found that the DEGs were enriched in 11 pathways, including the PI3K-Akt signaling pathway. Previous studies showed that several signal pathways could be involved in the progress and development of CRC [18-23]. PI3K/AKT is one of the classic pathways responsible for multiple biological processes, such as cell proliferation, differentiation, and migration. As a key effector of PI3K, Akt phosphorylation is closely related to CRC cell proliferation and apoptosis inhibition [18]. The specific inhibition of the PI3K/AKT signaling pathway could result in a decrease in proliferation and increase in apoptosis of the cells [24].

Furthermore, we constructed a PPI network and performed survival analysis along with verification of the RNA expression levels via the GEO database, DAVID and STRING online tools, Cytoscape software, and UALCAN online analysis. After the above series of operations, 4 hub genes, *CXCL2, SERPINE1, SPP1*, and *TIMP1*, were selected. Furthermore, we tested the expression of these 4 genes at the mRNA as well as protein levels and found that SPP1, TIMP1, and SERPINE1 had higher expression levels in colon adenocarcinoma cell lines than the colon epithelial cell line as well as in colorectal cancer tissue compared to para-cancerous tissue.

SPP1, named as osteopontin, is a phosphoprotein secreted by malignant cells. Cheng et al [25] found that SPP1 has a higher expression level in CRC tissues as compared to normal tissue, wherein its expression is closely associated with lymph node metastasis and the TNM stage. Inhibition of SPP1 in the HCT116 cells consequently resulted in the suppression of cellular growth, migration, and invasion. SPP1 overexpression could thus enhance the proliferation, migration, and invasion of CRC cells as well as the activation of the PI3K pathway, accompanied by the increased expression of Snail and MMP9 [26]. As one of the transcription factors of EMT, snail can inhibit the expression of E-cadherin, thereby promotes the activation of EMT in tumor cells [27]. Matrix metalloproteinase 9 (MMP9) is one of the members in the MMP family which is a class of zinc-dependent proteases with the main function of participating in the degradation of extracellular matrix. MMP9 is often overexpressed in malignant tumors and is associated with aggressive malignant phenotypes and poor prognosis in cancer patients [28]. Similarly, SPP1 could not only promote CRC cell growth and migration, but it also inhibited autophagy, which may be thus achieved by activating the p38MAPK signaling pathway [29]. In addition, a study by Choe et al. [30] also confirmed higher SPP1 expression along with shorter survival times in CRC patients.

TIMP1 is one of the members of the TIMP family which can bind to MMP substrates and mediate MMP degradation that is necessary for matrix renewal [31]. TIMP1 has been shown to be highly expressed in many types of cancer, consequently correlated to their poor prognosis [32-36]. TIMP1 may also promote liver metastasis in CRC [37]. After chemotherapy, high expression levels of TIMP1 in the serum have been closely related to the decline over survival (OS) [38]. Moreover, knockdown of TIMP1 leads to decreased proliferation of HT29 cells [39]. Huang [40], Zeng [41], and Zuo [42] et al. demonstrated that the expression levels of TIMP1 may be negatively correlated with the prognosis of COAD patients. Song et al. [43] found that TIMP1 knockdown could result in decreased proliferation, migration, and invasion as well as increased apoptosis *in vitro*, whereas there is weakened tumorigenesis and metastasis *in vivo* via the FAK-PI3K/Akt and MAPK pathways. The elevated TIMP1 levels in serum were related to lower tumor grade and shorter overall survival times of CRC patients [44].

SERPINE1, also called PAI-1, can regulate plasmin formation from plasminogen [45]. The expression level of SERPINE1 was seen to increase in CRC tissue and colon cancer cell lines showing active proliferation [46]. Moreover, overexpression of SERPINE1 led to decreased apoptosis and caspase-3 amidolytic activity in vascular smooth muscle cells (VSMC) compared to control [47]. LC-MS/MS analysis has shown that SERPINE1 may be a peripheral participant in the apoptosis of CRC cells [48]. Shun et al. found that the expression level of SEPINE1 was remarkably upregulated in tumor stroma and associated with the relapse of CRC stage II-III patients [49]. Several recent studies have suggested that SERPINE1 may be associated with prognosis of CRC patients [41, 50, 51].

CXCL2 belongs to the chemokine superfamily, and is responsible for the attraction of leukocytes to inflammatory sites apart from the regulation of angiogenesis, tumorigenesis [52]. Dietrich et al. found that there was significantly increased expression of CXCL2 in CRC, in contrast to normal tissue [53]. Further, there was increased expression of mRNA and protein levels of CXCL2 in colon cancer stem cells, whereas CXCL2 knockdown led to the decreased expression of EMT markers, MMPs, and Gai-2, which consequently promoted tumor initiation and development [54]. Down-regulation of CXCL2 destroyed the proliferation and colony formation in CRC cells [55]. Moreover, CXCL2 remarkably promoted the proliferation and migration of HUVECs [56]. But the qPCR results showed no significant difference in CXCL2 expression between HCT116 and NCM460 cell lines. This may be due to the source of the cell lines; more cell lines should thus be selected for verification in future studies.

In summary, in this study, we screened 323 DEGs from four COAD GEO datasets and identified *SERPINE1, SPP1*, and *TIMP1* as key genes which may be closely related to the prognosis of COAD in patients. Further investigations need to be performed to explore the mechanisms of these genes. These results would consequently provide candidate targets for the diagnosis and prognosis of CRC in patients, as well as new ideas for treatment.

## Author’s contributions

Yong Liu conceived the idea behind this paper; Chao Li and Xuewei Chen collected and analyzed the data; Lijin Dong and Ping Li drafted the paper; Rong Fan revised the final paper.

## Acknowledgements

This research was funded by the National Natural Science Foundation of China, No. 81201757 and 81973237; as well as the Basic research projects of Logistic University of Chinese People’s Armed Police Force, No.WHJ201702; and BWS17J025, AWS16J004. The visualization of GO analysis, including BP, MF and CC of DEGs, was performed via the online software ImageGP (http://www.ehbio.com/ImageGP/index.php/Home/Index/index).

## Ethics approval and consent to participate

Not applicable.

## Abbreviation

CRC: colorectal cancer;
GEO: Gene Expression Omnibus;
DEGs: differentially expressed genes;
DAVID: Database for Annotation, Visualization and Integrated Discovery;
GO: Gene ontology;
KEGG: Kyoto Encyclopedia of Gene and Genome;
BP: biological process;
CC: cellular component;
MF: molecular function;
STRING: Search Tool for the Retrieval of Interacting Genes/Proteins;
MCODE: Molecular Complex Detection;
TCGA: The Cancer Genome Atlas;
CXCL2: C-X-C Motif Chemokine Ligand 2;
SPP1: Secreted Phosphoprotein 1;
SERPINE1: Serpin Family E Member 1;
TIMP1: Tissue Inhibitor Of Metalloproteinases 1 ;
MAPK: Mitogen-Activated Protein Kinase;
PI3K: Phosphoinositide-3-Kinase;
HUVEC: human umbilical vein endothelial cells;
EMT: epithelial-mesenchymal transition;
MMPs: matrix metalloproteinases;
OS: over survival.

